# Discovery of malathion resistance QTL in *Drosophila melanogaster* using a bulked phenotyping approach

**DOI:** 10.1101/2022.07.19.500708

**Authors:** Stuart J Macdonald, Anthony D Long

## Abstract

*Drosophila melanogaster* has proven an effective system with which to understand the evolutionary genetics and molecular mechanisms of insecticide resistance. Insecticide use has left signatures of selection in the fly genome, and both functional and quantitative genetics studies in the system have identified genes and variants associated with resistance. Here, we use *D. melanogaster* and leverage a bulk phenotyping and pooled sequencing “extreme QTL” approach to genetically dissect variation in resistance to malathion, an organophosphate insecticide. We resolve two QTL (Quantitative Trait Loci), one of which implicates allelic variation at the cytochrome P450 gene *Cyp6g1*, a strong candidate based on previous work. The second shows no overlap with hits from a previous genomewide association study (GWAS) for malathion resistance, recapitulating other studies showing that different strategies for complex trait dissection in flies can yield apparently different architectures. Notably, we see no genetic signal at the *Ace* gene. *Ace* encodes the target of organophosphate insecticide inhibition, and GWAS have identified strong *Ace*-linked associations with resistance. The absence of QTL implicating *Ace* here is most likely because our mapping population does not segregate for several of the known functional polymorphisms impacting resistance at *Ace*, perhaps because our population is derived from flies collected prior to the widespread use of organophosphate insecticides. Our fundamental approach can be an efficient, powerful strategy to dissect genetic variation in resistance traits. Nonetheless, studies seeking to interrogate contemporary insecticide resistance variation may benefit from deriving mapping populations from more recently collected strains.

## INTRODUCTION

Increased agricultural productivity over the last century has been, at least in part, promoted by broader use of insecticides to protect crops agains insect pests (Tudi et al. 2021). Equally, insecticides are widely-employed to protect humans against insect vectors of infectious disease (World Health Organization 2006). A side-effect of this widespread insecticide use is that populations of many insect species – those that are the targets of insecticides, as well as non-target species – have evolved strong resistance to important classes of insecticides (e.g., Moyes et al. 2017; Hemingway and Ranson 2000; Rivero et al. 2010).

A route to understanding insecticide resistance mechanisms is to resolve the variants that have been selected in populations (Hawkins et al. 2018). Critical insight into the genetic evolution of resistance has come from direct work with a range of pest insect/insect vector systems, including various *Anopheles/Culex* mosquito species (see Hemingway et al. 2004), and *Lucilia* blowflies (e.g., Hartley et al. 2006). Nonetheless, despite not itself being either a disease vector or a crop pest (although see Hall et al. 2018) *Drosophila melanogaster* has emerged as a useful and practical system with which to study insecticide resistance (Scott and Buchon 2019; Ffrench-Constant et al. 2004). This is due to the relative ease with which *D. melanogaster* can be genetically manipulated to test the molecular genetic role of genes/variants, the availability of a huge array of genomescale data on sequence/transcriptome variation in *D. melanogaster*, and the existance of living resources enabling the genetic dissection of trait variation (King et al. 2012a; King et al. 2012b; Huang et al. 2014; Mackay et al. 2012). Additionally, studies have demonstrated that natural populations of *D. melanogaster* have been impacted by insecticide use, with signatures of adaptation around genes known to be involved in the response to insecticide exposure (Duneau et al. 2018; Garud et al. 2015; Karasov et al. 2010)

Several studies have sought to use the DGRP (*Drosophila* Genetic Reference Panel), a panel of ~200 sequenced inbred strains (Huang et al. 2014; Mackay et al. 2012), to resolve the genetic archictecture of insecticide resistance in this lab-derived *D. melanogaster* population using a GWAS (Genomewide Association Study) design. In some cases studies have recovered GWAS hits at known insecticide targets, such as the *Ace* (*Acetylcholine esterase*, FBgn0000024) gene (Battlay et al. 2018; Duneau et al. 2018) which can be inhibited by organophosphate and carbamate insecticides. Additionally, GWAS studies have implicated variation at genes that encode cytochrome P450 detoxification enzymes, such as *Cyp6g1*/FBgn0025454 (Battlay et al. 2018; Battlay et al. 2016) and *Cyp6a23*/FBgn0033978 (Battlay et al. 2018; Duneau et al. 2018) in insecticide resistance in flies.

The present study was motivated by two observations. First, genetic dissection regimes relying on panels of inbred fly strains can be quite inefficient to execute; It is a technical challenge to accurately measure a phenotype on multiple individuals, from both sexes, from dozens to hundreds of different strains. Additionally, such designs are subject to confounding microenvironmental noise, since different genotypes are reared/assayed in different vials/bottles. Approaches that allow test individuals to be communally reared/assayed under common garden conditions, and enable bulked phenotyping can mitigate these issues.

Second, for a given phenotype, the genetic architecture uncovered can depend on the genetic dissection approach employed. In addition to the DGRP, a number of studies have used the *Drosophila* Synthetic Population Resource (King et al. 2012a; King et al. 2012b) to dissect trait variation. The DSPR consists of two sets of genotyped Recombinant Inbred Lines (RILs), each derived from an advanced generation intercross initiated with 8 inbred founder strains, that enable QTL (Quantitative Trait Locus) mapping (Long et al. 2014). For two virus infection phenotypes, studies in both the DGRP (Magwire et al. 2012) and DSPR (Cogni et al. 2016) have strongly implicated variation at the same loci (the *pst*/FBgn0035770 gene for resistance to Drosophila C Virus infection, and the *ref(2)P*/FBgn0003231 gene for resistance to the sigma virus DMelSV). However, dissection of three xenobiotic and stress resistance phenotypes in both the DSPR and DGRP revealed distinct architectures (Najarro et al. 2017; Najarro et al. 2015; Everman et al. 2019); the DSPR suggesting a significant fraction of trait heritability is encapsulated by a handful of mapped loci, whereas in the DGRP few variants yield hits passing a strict, genomewide threshold, and even sub-threshold associations are not enriched within the QTL intervals resolved by the DSPR. Since studies examining variation in insecticide resistance have emphasized the DGRP system (Battlay et al. 2018; Battlay et al. 2016; Green et al. 2019; Schmidt et al. 2017; Denecke et al. 2017; Duneau et al. 2018), an investigation of the genetic archictecture of insecticide resistance that derives from other approaches is of interest.

In the present study we dissect variation in resistance to malathion, an organophosphate insecticide, using an “extreme QTL” or X-QTL mapping design (Ehrenreich et al. 2010). We employ bulked phenotyping and pooled sequencing of malathion-selected and control individuals derived from a population constructed by mixing several hundred DSPR RILs. Our use of this design in a previous study of caffeine resistance recapitulated QTL isolated via traditional RIL-by-RIL phenotyping (Macdonald et al. 2022; Najarro et al. 2015), while being – in our opinion – more efficient to execute. Since variation in malathion resistance has been examined previously using the DGRP – Battlay et al. (2018) found very strong associations at *Ace* and a more modest association at *Cyp6g1* – we could execute a direct comparison of the outcomes of two different approaches to genetic analysis in flies.

## MATERIALS & METHODS

### Fly population

The X-QTL base population used here is a derivation of the one created by Macdonald et al. (2022), and consists of the outbred descendants of a mix of RILs from the DSPR collection (King et al. 2012a; King et al. 2012b).

Briefly, 8 highly-inbred strains (Chakraborty et al. 2019; King et al. 2012b) were used to found a synthetic population that was maintained at large population size for 50 generations. Subsequently a large set of 8-way, advanced generation RILs were developed via 25 generations of sibling mating (yielding the DSPR lines). The X-QTL base population was created by collecting 10 embryos from each of 663 DSPR pA (“panel A”) RILs, and releasing emerged adults into a 1 cubic foot population cage. The population was maintained in this cage for 21 non-overlapping generations in an incubator (25°C, 50% relative humidity, 12-hour light / 12-hour dark), replacing the 9-12 rearing bottles in the cage approximately every 2 weeks. Following this period flies were moved to an 8 cubic foot population cage, the cage was moved out of the incubator (so was subject to a more variable environment), and each week 3 of the 12 rearing bottles in the cage were switched for fresh bottles (so the population now experienced overlapping generations). Assuming that the population experienced 1 generation every 2 weeks with this maintenance regime, eggs were collected from the population ~28 (experimental replicate 1) and ~48 (replicate 2) generations after its founding from DSPR RILs. Adults emerging from these eggs were used for the bulk malathion resistance assay described below.

### Rearing experimental animals

For each experimental replicate, test animals were derived from the base population as follows, with additional detail in Macdonald et al. (2022). We added five 100-mm diameter petri dishes containing apple juice agar and a small dab of live yeast paste to the base population cage overnight. The next day eggs were removed from plates, suspended in 1X PBS (phosphate-buffered saline), rinsed with additional PBS, and 12-μl of eggs were pipetted into standard *Drosophila* rearing vials (Fisher Scientific, AS515) each containing ~10-ml of cornmeal-yeast-molasses media. Egg pipetting in this fashion has been shown to yield relatively homogeneous egg density over vials (Clancy and Kennington 2001). Two days following the first emergence of adults, all emerged flies were moved to fresh media vials. The next day, we sexed flies over CO2 anesthesia, collecting a series of single-sex groups of 50 flies into fresh media vials. Flies – then 3-5 days old – were assayed the following day. All rearing, maintenance and testing (below) of experimental flies was conducted at 25°C, 50% relative humidity, and on a 12-hour light / 12-hour dark cycle.

### Collecting control, unselected animals

From each single-sex vial of experimental flies we aspirated 4 flies prior to the malathion resistance assay. These arbitrarily-collected control flies represent a sample of the base population allele frequency. We collected a total of 120 control males and females for replicate 1, although only a subset (65 and 43, respectively) were employed for pooled DNA isolation in order to match the number of malathion-selected animals obtained (below). We collected 264 control flies of each sex for replicate 2, all of which were used for pooled DNA isolation.

### Malathion resistance assay

The design of the assay was largely copied from previous studies of insecticide resistance in flies (e.g., Battlay et al. 2018; Schmidt et al. 2017; Schmidt et al. 2010). Malathion (Millipore-Sigma, 36143-100MG) was added to acetone (Fisher Chemical, A929) at a concentration of 2μg/ml, and 500-μl of the mix was added to a series of 20-ml glass scintillation vials (Fisher Scientific, 03-337-5). The interior walls of these vials were coated in malathion by rolling on a hotdog warmer (Grand Slam, HDRG12) with the heat off for ~15-min in a fume hood to evaporate the acetone. Malathion-coated scintillation vials were left overnight at room temperature before being used for the assay.

Test flies were tipped from single-sex holding vials to malathion-containing vials, and these were plugged with 1/2 of a large cotton ball (VWR, 14224-518) dampened with 1-ml of 10% sucrose solution (Fisher Chemical, S5). Assays were initiated within the first (replicate 1) or second (replicate 2) hour following lights on. After a period of exposure (210-245 minutes for replicate 1, 190-230 minutes for replicate 2), flies – both living and dead – were transferred from malathion vials to normal media vials and left for 24 hours. This was done because pilot experiments indicated that a fraction of the animals remaining alive immediately following the assay would succumb to the effects of the insecticide after several hours. Plus, discrimating live from dead animals was considerably easier after this period.

### Collecting malathion-selected animals

One day following the assay, alive and dead animals were separated over CO_2_ and counted. In replicate 1, 65 of 1,303 males (5.0%) and 43 of 1,338 females (3.2%) survived, while in replicate 2, 403 of 2,881 males (14.0%) and 693 of 2,817 females (24.6%) survived. Given simulation data presented in Macdonald et al. (2022), the scale of the experiment was lower than optimal in replicate 1, while the intensity of the selection was sub-optimal for replicate 2.

### DNA isolation, library construction, and sequencing

We isolated DNA from each pool of animals (2 replicates × 2 treatments × 2 sexes = 8 total pools) via the Gentra Puregene Cell Kit (Qiagen, 158767) using straightforward extensions of the manufacturer’s protocol, and the resulting DNA was quantified using a fluorometer (Qubit dsDNA BR Assay Kit, ThermoFisher, Q32853). Subsequently, we used 400-ng of each DNA sample to construct indexed sequencing libraries (Illumina DNA Prep Tagmentation, 20018705; Illumina Nextera DNA CD Indexes, 20018708). Libraries were mixed by replicate into two 4-plexes, and sequenced on an Illumina NextSeq 550 instrument. We obtained 21.2-26.0 million PE150 read pairs for each replicate 1 sample, and 31.2-37.7 million PE75 for each replicate 2 sample.

### Read mapping, SNP calling, and haplotype frequency estimation

For each X-QTL sample, along with the set of 8 inbred strains that founded the pA DSPR population (King et al. 2012b), raw reads were first mapped to the *D. melanogaster* reference genome (Release 6, dm6) via bwa-mem (Li 2013). This resulted in 50X coverage for the replicate 1 female pools, and 32-39X coverage for the remaining pools. Previous work indicates coverage at this level enables robust haplotype frequency estimation (Macdonald et al. 2022). Next, the bcftools mpileup, call and query commands (Li 2011) were employed to generate a file of REF and ALT counts at all SNPs for each sample (founders plus X-QTL samples), and this was converted to REF allele frequencies per sample per SNP (often 0/1 for the inbred founders).

We call haplotypes for each X-QTL sample in windows of 1.5-cM, stepping through the genome in 0.05-cM increments, using a procedure described in more detail previously (Linder et al. 2020; Macdonald et al. 2022). Briefly, for each X-QTL pooled sample, and within each window, we use the R/limSolve package (Soetaert et al. 2009) to find the set of 8 proportions (summing to 1) that minimizes the sum of the weighted squared differences between the known founder haplotypes, and the observed frequency of each SNP in the window in that sample. Experimental validation of the accuracy of this approach in an 18-way yeast population, and an 8-way DSPR-derived population is presented elsewhere (Linder et al. 2020; Macdonald et al. 2022).

### X-QTL genome scan

Mapping is performed by executing a statistical test at each window (above) along the genome. First, the set of 8 inferred founder haplotype (*H*) frequencies from each replicate (*R*), and from the control and malathion-selected treatments (*T*) are arcsine square-root transformed (*ASF*). We chose to treat the male and female tests within each experimental replicate as independent, so *R* = 4 (2 experimental replicates × 2 sexes), and we are therefore geared to identify effects that are consistent in each sex. We then test for differentiation between treatments using the ANOVA *ASF* ~ *H* + *TRT* + *H*×*TRT*, testing for the effect of the *H*×*TRT* interaction using *R*×*H*×*TRT* as the error term, returning –log_10_(*P*) values. Previous simulation work over a broad parameter space indicates that –log_10_(*P*)=4 holds the QTL false positive rate roughly at ~5% genomewide (Linder et al. 2020; Macdonald et al. 2022). However, we acknowledge the present study is at the lower end of the factors that impact power of the DSPR-based X-QTL design (Macdonald et al. 2022). Finally, following smoothing of the –log_10_(*P*) values across each chromosome (via LOESS, to accommodate window-to-window variation in the test statistic), we called QTL peaks, and automatically extracted 3 –log_10_(*P*) drop (“LOD drop”) confidence intervals, which in simulations encapsulate the true position of the causative locus ~95% of the time (Macdonald et al. 2022).

### Gene functional annotations

We used FlyBase (version FB2022_03, Gramates et al. 2022) to identify plausible candidate genes within mapped QTL, marking genes if they were tagged with the Gene Ontology terms “detoxification” (GO:0098754) or “response to insecticide” (GO:0017085), or if they are members of the following FlyBase Gene Groups – all known players in the detoxification pathway (Xu et al. 2005); cytochrome P450 genes (FBgg0001222), glutathione s-transferases (FBgg0000077), other carboxylesterases (FBgg0001375), GT1 family of UDP-glycosyltransferases (FBgg0000797), or ATP-binding cassette transporters (FBgg0000547).

### Testing for a heritable effect of malathion selection

Successful selection for malathion resistance in a population should result in the progeny of the selected population showing increased resistance. For experimental replicate 1, prior to freezing animals for subsequent DNA isolation, we allowed the control animals (120 males and 120 females) and the malathion-selected animals (65 males and 43 females) to lay eggs in regular media vials for ~24-hours. In the following generation, we transferred mixed-sex groups of progeny adults to fresh vials 2 days following the first adult emergence, left flies to mate/age for 48 hours, then collected 20 vials of 10 female progeny from both the control and selected populations over CO_2_ anesthesia. The following day (at around 1-hour following lights on), when progeny animals were 3-5 days old, all 40 vials were tipped into malathion exposure vials (described above) and the number of dead flies were manually counted by the same investigator periodically over the next 9-10 hours. Nearly all flies died during this period.

### Data availability

Raw malathion X-QTL sequencing data generated for this project is available on the NCBI SRA under BioProject Accession PRJNA857080, while the DSPR founder FASTQs required for analysis are available via Accession SRP011971. Scripts to run the analyses presented are available on FigShare (INSERT_URL_HERE).

## RESULTS & DISCUSSION

We sought to employ a bulk phenotyping/genotyping X-QTL strategy to resolve genomic regions contributing to resistance to the insectide malathion in *D. melanogaster*. The base population for selection was generated by mixing several hundred strains from the DSPR collection (King et al. 2012a; King et al. 2012b), resulting in an outbred, highly-recombinant population segregating for at most 8 haplotypes at any given position. Subsequently, our experiment followed the same fundamental design we have employed to isolate caffeine resistance QTL (Macdonald et al. 2022). Over 2 experimental replicates, samples of male and female flies from the base population were exposed to malathion, and after a period of exposure surviving animals were retained. Each pool of malathion-selected animals, along with matching pools of unselected, control animals sampled randomly from the pre-exposure cohorts of experimental animals, were subjected to bulk DNA isolation and sequencing library construction (2 replicates × 2 sexes × 2 treatments = 8 X-QTL samples in total). Following sequencing, we inferred the frequencies of the 8 possible haplotypes for each of the pooled X-QTL samples at intervals across the genome. Finally, we executed a test to identify consistent differentiation between control and selected samples at each position, resolving locations – QTL – showing significant allele-frequency shifts.

### Selection results in progeny with greater malathion resistance

Prior to freezing off selected and control animals for experimental replicate 1, the two cohorts were allowed to lay eggs, and their adult female progeny tested for malathion resistance. In the ~10-hours over which flies were monitored, 199/200 (99.5%) control female progeny and 183/195 (93.8%) selected female progeny died. Even ignoring these “censored” individuals, progeny of malathion-selected cohorts live substantially longer than progeny of control animals (**Figure 1**); the control and selected means are 185, and 286-minutes, respectively (Welch’s *t*-test = 9.98, *p* < 10^-16^). Malathion-selected pools of individuals are enriched for alleles that confer greater resistance to the toxic effects of the insecticide.

**Figure 1.**
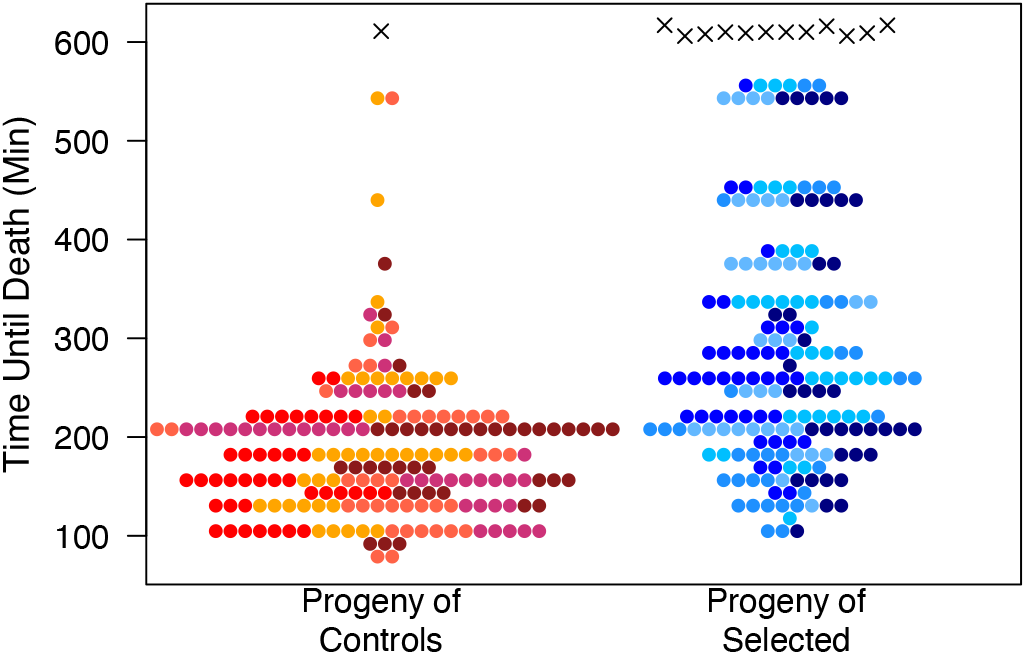
Female progeny of malathion-selected animals are more resistant to the insecticide than progeny of controls. Around 200 female progeny of the replicate 1 cohorts of selected/control animals were assayed for malathion resistance, and the number of dead animals was counted periodically over a ~10-hour exposure period. Each point represents a single female. Animals dying during the exposure period (colored, filled circles) are assumed to have died at the midpoint between sequential counting times. Colors represent the 5 rearing vials of origin for each sample of test flies. Those animals dying after the exposure period (crosses) are presented at the last time they were scored as alive. The progeny of selected females are significantly more resistant to malathion (*t*-test, *p* < 10^-16^).

### Two mapped autosomal loci contribute to malathion resistance in the DSPR

We considered the male and female tests within each experimental replicate as independent, yielding 4 control-selected pairs of samples. We did this – rather than considering sexes separately – since we are likely underpowered to identify sex-specific effects given our limited level of replication (Macdonald et al. 2022). Pooling across sexes is supported by the result that males and females from the same set of 170 DGRP strains have strongly correlated malathion resistance phenotypes (Pearson’s *r* of 0.81-0.89), and by association mapping that appears to implicate the same major malathion resistance loci in both sexes (Battlay et al. 2018).

An X-QTL genome scan contrasting control and selected population frequencies was executed across the genome to identify consistent haplotype frequency shifts due to selection, and analysis revealed 2 peaks rising above a –log_10_(*P*)=4 threshold, one on each autosome **(Figure 2**). Confidence intervals on QTL locations implicate fairly wide intervals, and hundreds of protein-coding genes (**Table 1**), likely due to the limited replication in our experiment (see Macdonald et al. 2022).

**Figure 2.**
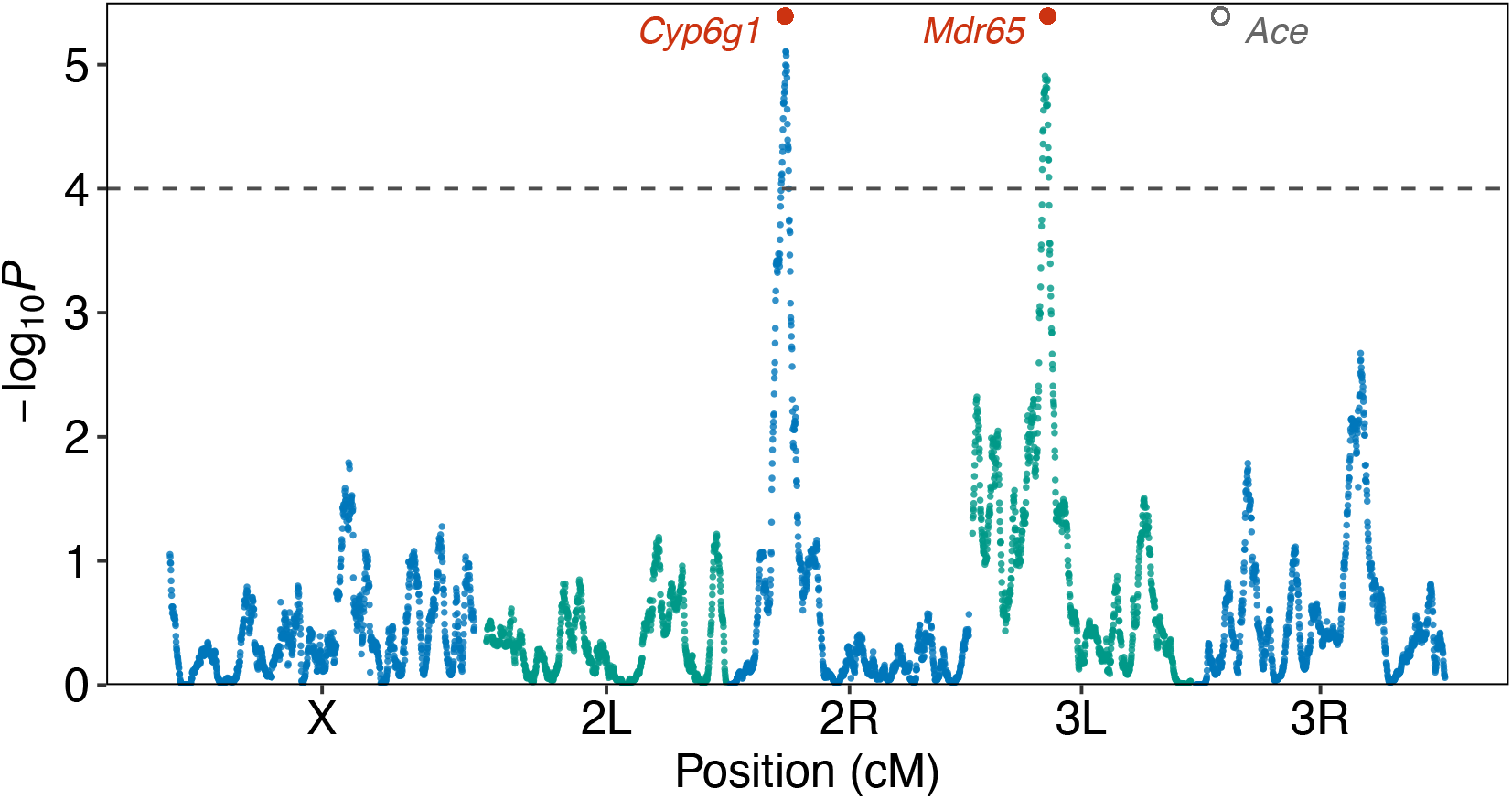
Two QTL for malathion resistance in the DSPR. The –log_10_(*P*) value is the result of contrasting haplotype frequencies of all pairs of control and selected populations in a series of 1.5-cM windows along the genome. Peaks surviving a –log_10_(*P*)=4 threshold are apparent on chromosome arms 2R and 3L. The locus on 2R includes the well-known *Cyp6g1* insecticide resistance candidate gene, which was previously implicated in natural malathion resistance by a GWAS (Battlay et al. 2018). An ABC transporter gene *Mdr65* that impacts insecticide resistance when knocked out/down (Denecke et al. 2017; Sun et al. 2017) is within the 3L interval. There is no indication that the *Ace* gene – a target of inhibition by organophosphates such as malathion, known to segregate for functional variation impacting insecticide resistance in flies, and a major GWAS hit in a previous study of malathion resistance (Battlay et al. 2018) – is associated with phenotype in our DSPR-based X-QTL mapping study; The maximum –log_10_(*P*) score in a 2-Mb window centered on *Ace* is 0.83.

**Table 1.** Mapped malathion resistance QTL.

| Chr | Peak Position (bp)<sup>a</sup> | Physical Interval (bp)<sup>a, b</sup> | QTL size<sup>b</sup> |  |  |
| --- | --- | --- | --- | --- | --- |
|  |  |  | Mb<sup>a</sup> | cM | Genes<sup>c</sup> |
| Chr2R | 12,294,089 | 10,966,645-13,213,848 | 2.25 | 4.7 | 344 |
| Chr3L | 6,162,469 | 5,515,636-6,735,645 | 1.22 | 3.8 | 145 |

<sup>a</sup> Based on Release 6 (dm6) of the *D. melanogaster* reference genome.
<sup>b</sup> Intervals are defined as 3 $-\log_{10}(P)$ drops from the QTL peak.
<sup>c</sup> Specifically the number of protein-coding genes in the interval. See **Supplementary Table 1** for additional details on these genes.

A feature of any mapping system based on a multiparent cross is that the parental haplotypes at a mapped locus can be interrogated for whether they posses phenotype increasing/decreasing alleles. **Figure 3** shows that the Chr2R QTL is “driven” by resistance alleles harbored by founders A5 and A6 (these alleles increase in frequency in the selected samples), while A7 appears to confer susceptibility (and the allele decreases in frequency in the selected samples). Similarly, at the Chr3L locus it appears that the AB8 founder allele confers greater resistance, while A2/A3 reduce resistance. Notably, examination of the haplotype frequency differences between each matched pair of control and selected pools (**Figure 4**) suggests there is a general concordance between frequencies in males and females, suggesting that our analysis of the data without regard to sex was reasonable.

**Figure 3.**
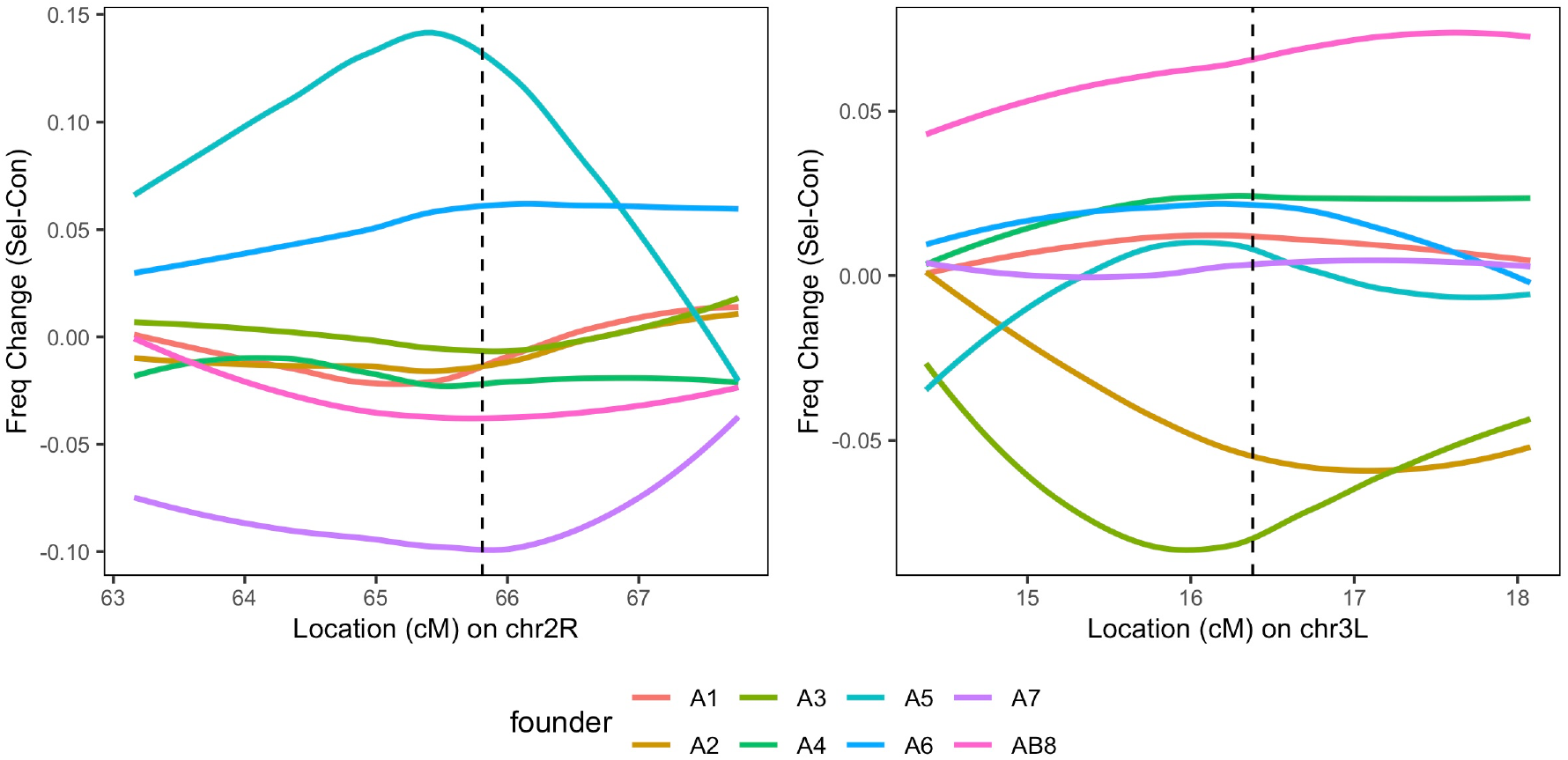
Haplotype frequency differences between control and selected groups at mapped X-QTL. Each plot shows the frequency difference (selected minus control) for each of the 8 founder haplotypes through the 3 –log_10_(*P*) drop interval implicated by each QTL (*left* = Chr2R, *right* = Chr3L), with QTL peaks indicated by vertical dashed lines. Values above zero indicate the haplotype frequency is greater in the selected group.

**Figure 4.**
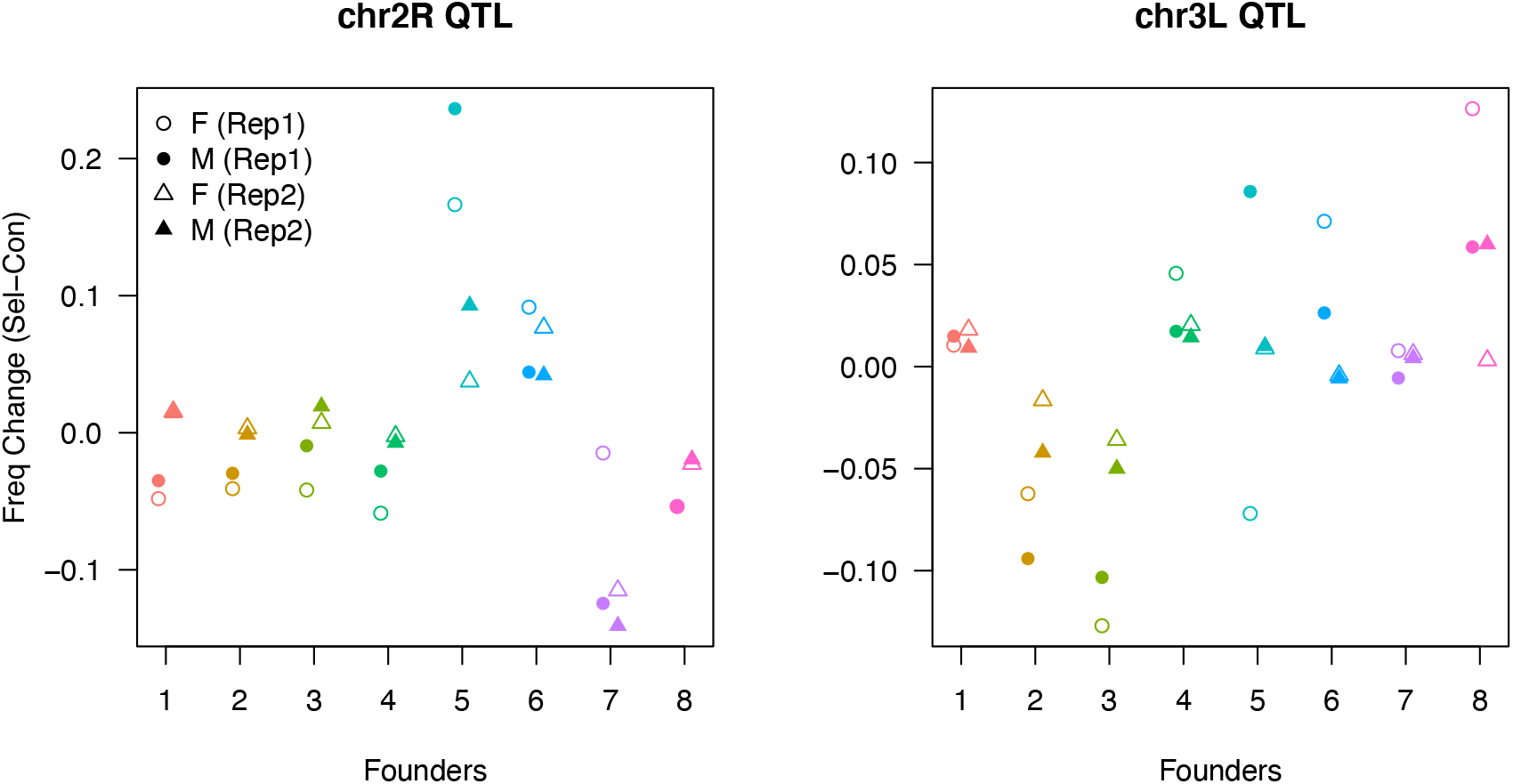
Consistent haplotype frequencies over experimental replicates/sexes at mapped QTL peaks. Each plot shows the frequency difference (selected minus control) for each of the 8 DSPR pA founder haplotypes (1 -7 = A1 -A7, 8 = AB8) at the two X-QTL peaks, separately for each experimental replicate (Rep1, Rep2) and sex (F, Females; M, Males).

### A well-known insecticide resistance gene, *Cyp6g1*, resides within the Chr2R mapped locus

Both of our X-QTL intervals encompass fairly large numbers of genes (**Table 1**, **Supplementary Table 1**), so we attempted to identify plausible candidate genes by marking those that fall into any of 7 formally-defined functional categories relevant to detoxification and insecticide resistance (see Materials & Methods). We also made use of results from Salces-Ortiz et al. (2020, see their supplementary table s1) to highlight genes that show differential expression in adult female gut tissue in response to malathion.

For the Chr2R QTL, 9 genes have annotations suggesting a role in detoxification and/or insecticide resistance (**Table 2**), and of these 7 were shown to be differentially-expressed in response to malathion (Salces-Ortiz et al. 2020). Cytochrome P450 gene *Cyp6g1* emerges as the strongest candidate to underlie variation at this locus. The gene has a well-defined role in resistance to DDT and other insecticides (Daborn et al. 2001; Daborn et al. 2002; Schmidt et al. 2010; Harrop et al. 2014; Chung et al. 2007), population-based GWAS have identified hits in/near *Cyp6g1* for malathion (Battlay et al. 2018) and azinphos-methyl resistance (Battlay et al. 2016), and a *Cyp6g1* knockout reduces resistance to malathion (Battlay et al. 2018). Additionally, A6 – one of the two founders conferring resistance at this QTL (**Figure 3**) – possesses 2 copies of *Cyp6g1*, one with a full-length, and one with a fragment of an *Accord* transposable element just upstream of the start of the gene copy (Chakraborty et al. 2019; DSPR Variant UCSC Browser 2019). Previous work has indicated an *Accord* insertion in this region increases *Cyp6g1* transcription (Daborn et al. 2002; Chung et al. 2007). Indeed, individuals carrying the A6 haplotype at *Cyp6g1* exhibit higher gene expression in adult female heads than those carrying other DSPR founder haplotypes at this position (King et al. 2014; Chakraborty et al. 2019). Interestingly, founder A5, which shows the largest increase in frequency with selection at the Chr2R QTL (**Figure 3**, **Figure 4**), possesses only a single copy of *Cyp6g1* and no structural variation is evident at the gene (Chakraborty et al. 2019; DSPR Variant UCSC Browser 2019). Clearly however, other genetic factors that are less pronounced than structural variants can contribute to phenotype, and could be present in the A5 *Cyp6g1* region.

**Table 2.** Plausible candidate genes present within QTL intervals.

| QTL | Gene Symbol<sup>a</sup> | Functional Evidence<sup>b</sup> | Malathion DE Response<sup>c</sup> |
| --- | --- | --- | --- |
| Chr2R | *Cyp12d1-d*<sup>d</sup> | P450, Insecticide | B+ / C+ |
|  | *Cyp12d1-p*<sup>d</sup> | P450, Insecticide | B+ / C+ |
|  | *Sod3* | Detox | ns |
|  | *Cyp6g1* | P450, Detox, Insecticide | B+ / D– |
|  | *Cyp6g2* | P450, Detox, Insecticide | D– |
|  | *Cyp6t3* | P450, Detox, Insecticide | ns |
|  | *Cyp301a1* | P450 | C– / D– |
|  | *Cyp9h1* | P450 | C+ |
|  | *Mdr49* | ABC, Insecticide | B+ / D– |
| Chr3L | *Spo* | P450 | D+ |
|  | *CG10226* | ABC | B+ / C+ |
|  | *Mdr65* | ABC, Insecticide | ns |
|  | *sfl* | Est | ns |

<sup>a</sup> Full detail on the genes can be found in **Supplementary Table 1**.
<sup>b</sup> Functional evidence associated with each gene on FlyBase (Gramates et al. 2022) is encoded as follows (see Materials & Methods): P450 = cytochrome P450, Est = carboxylesterase, ABC = ABC transporter, Detox = gene involved in detoxification, Insecticide = gene has a role in the response to insecticide.
<sup>c</sup> Salces-Ortiz et al. (2020) identified genes as being upregulated (+) or downregulated (–) in adult female gut tissue in response to malathion in each of 4 independent *D. melanogaster* strains (encoded here by the letters A-D). See legend of **Supplementary Table 1** for more detail.
<sup>d</sup> The reference *D. melanogaster* genome harbors *Cyp12d1-d* and *Cyp12d1-p*. These are nearly identical copies of the same gene that is subject to copy number variation in natural populations (Najarro et al. 2015; Good et al. 2014; McDonnell et al. 2012; Schrider et al. 2013; Rivero et al. 2010).

The cytochrome P450 gene *Cyp12d1* is also present within the Chr2R QTL interval (**Table 2**). This is notable since transgenic overexpression of this gene results in greater malathion resistance (Battlay et al. 2018). The DSPR segregates for copy number variation at *Cyp12d1*, and of the DSPR pA founders, only founder A7 harbors 2 copies (Najarro et al. 2015). However, at the Chr2R QTL this founder appears to be associated with reduced resistance (**Figure 3**). Coupled with the observation that *Cyp12d1* copy number is not associated with resistance towards a different insecticide, DDT (Schmidt et al. 2010), *Cyp12d1* seems to represent a less compelling candidate to harbor functional variation yielding the Chr2R malathion resistance QTL we map. Ultimately, our work appears to recapitulate the GWAS result showing that variation at *Cyp6g1* impacts malation resistance (Battlay et al. 2018).

### A new malathion resistance locus

The malathion resistance GWAS of Battlay et al. (2018) revealed a set of 273 variants (of the >1.8 million tested) that survived a genomewide significance threshold (1.25 × 10^-6^ – the nominal 10^-5^ threshold that is commonly used in DGRP publications (Mackay and Huang 2018) corrected for the 4 GWAS executed on different malathion phenotypes in each sex). None of these variants are within our Chr3L QTL. One variant showed an association with a single resistance phenotype in males that survived a nominal *P*<10^-4^ threshold. This site is located in the *Lkr*/FBgn0035610 gene that encodes a G-protein coupled receptor involved in the regulation of feeding (Al-Anzi et al. 2010). Further work would be needed to evaluate whether variation at *Lkr* plays a role in malathion resistance.

Four genes at the Chr3L locus have existing annotations suggesting a role in resistance to toxicants (**Table 2**), two of which show upregulated expression in response to malathion exposure (see Salces-Ortiz et al. 2020). Despite the lack of an observed expression change in response to malathion, *Mdr65*/FBgn0004513 is perhaps the most likely of the candidates to segregate for insecticide-relevant variation; Previous work has shown that RNAi and null mutations at the gene result in reduced malathion resistance (Sun et al. 2017), and CRISPR-based *Mdr65* knockouts yield lower resistance to a range of insecticides (Denecke et al. 2017), although malathion was not among the panel of insecticides tested. Examination of the *Mdr65* gene sequence among the DSPR pA founder chromosomes does not reveal any notable structural variation (DSPR Variant UCSC Browser 2019). Nonetheless, that our study has inferred “high” and “low” alleles at this QTL (**Figure 3**) could enable explicit tests of whether *Mdr65* confers this variation, for instance by exploiting reciprocal hemizygosity testing (Stern 2014; Steinmetz et al. 2002) or allele swaps (Lamb et al. 2017).

### No signal of *Ace*-associated malathion resistance variation in the DSPR

Malathion is an acetylcholinesterase inhibitor. Five naturally-occuring amino acid changes – F115S, I199V, G303A, F368Y, G406A – in the *D. melanogaster Ace* gene have been shown in functional assays to confer some resistance to the toxic effects of insecticide exposure (Menozzi et al. 2004; Mutero et al. 1994; Shi et al. 2004). Three of these – I199V, G303A, F368Y – segregate in the DGRP, and all three are among the set of 62 variants shown in the Battlay et al. (2018) GWAS to survive a 3.33 × 10^-9^ threshold for at least one combination of malathion phenotype and sex. Indeed, of these 62 hits, 8 are within the 36.6-kb of the genome spanned by the *Ace* gene, and 54 (87%) reside in a 135.4-kb genomic interval that includes *Ace*. Additionally, hits at *Ace* were identified in a GWAS for resistance to parathion (Duneau et al. 2018), another organophosphate insecticide.

Despite the evidence of the role of *Ace* in insecticide resistance, we do not map a QTL at the *Ace* gene in our study, and all –log_10_(*P*) values in the region are well below our genomewide statistical threshold (**Figure 2**). Examining the DSPR founder sequences (King et al. 2012b; Chakraborty et al. 2019) reveals that none of the five functional changes listed above are present. This is perhaps because all the DSPR founders are derived from flies collected from nature in the mid-1950’s to the late 1960’s, right around the period when organophosphate insecticides started to be deployed widely (Casida and Quistad 1998). Thus, while we clearly lack power to find all variants associated with phenotype (Macdonald et al. 2022), and there will certainly be lab-to-lab variation in the resistance assay employed, since previous studies have identified massive effects on organophosphate resistance at variants in/near *Ace* (Battlay et al. 2018; Duneau et al. 2018), the absence of QTL implicating *Ace* in our study is most likely due to limited *Ace*-linked, insecticide-relevant variation present in our mapping panel.

### Differences in the inferred archictecture of trait variation driven by the genetic analysis strategy employed

Similar to our previous studies with the DGRP and DSPR panels of inbred strains (Everman et al. 2019; Najarro et al. 2017; Najarro et al. 2015), the present study again shows that genetic dissection of a given trait with different approaches can reveal a distinct underlying architecture. We appear to replicate the previously-identified effect on malathion resistance of natural allelic variation at *Cyp6g1*, but fail to repeat the hit at *Ace*, most likely because our base population does not segregate for strong-effect functional variants at the gene.

We additionally identify a QTL at a position in the genome where no genomewide significant malathion resistance hits were discovered in the DGRP (Battlay et al. 2018). This could be because the variant(s) giving rise to this QTL are absent in the DGRP; Indeed, only a fraction of the segregating variation in the two panels is shared (King and Long 2017). Alternatively, the causative variant(s) may be at very low frequency in natural populations. This would render them challenging to identify using any population-based GWAS approach (Spencer et al. 2009), but if captured in the founders, would be amenable to discovery in a multiparental advanced intercross design like the DSPR (King et al. 2012a). Finally, the variant(s) could simply have modest effects. With just 200 inbred lines, power to identify a site explaining 4% of the genetic variation for a trait in the DGRP is below 10% (Mackay and Huang 2018), so power deficits could explain the absence of a DGRP GWAS hit within the region of our Chr3L locus. Exact reasons aside, our work shows that different genetic dissection study designs can provide complementary insight into trait variation.

### An X-QTL approach is practical and efficient for genetic dissection of resistance traits

The present study did not fulfill all criteria enabling the highest resolution, highest power X-QTL study; Ideally we would have executed selection on larger cohorts of individuals, applied stronger selection while simultaneously retaining larger pools of selected animals, and repeated the experiment several more times (see Macdonald et al. 2022 for optimal parameters based on simulations). Nonetheless, similar to our previous study (Macdonald et al. 2022), we succeeded in replicating loci previously identified for our target xenobiotic resistance trait. The X-QTL approach is fairly efficient in terms of personnel time, and is particularly effective for stress/toxicant resistance phenotypes; it is generally straightforward to conceive of bulked phenotyping regimes for such traits, and the experimental animals “self sort” into the selected cohort – resistant animals remain alive at some time following a challenge.

Of course, the X-QTL approach will not be appropriate for all labs, or all questions. Unlike assaying a trait in a series of stable, genotyped strains, pooled sequencing is necessary. That said, with the necessary coverage per pool on the order of 30-50X (above and Macdonald et al. 2022), this can be cost-effective for *D. melanogaster* with current short-read sequencing technologies. X-QTL mapping also does not result in individual-level genotypes or phenotypes. This means one cannot take advantage of other phenotypes measured on the same genotypes to more directly explore connections among traits (the DGRP and DSPR panels have both been examined for many organismal and molecular phenotypes). It also renders X-QTL unable to dissect the contribution of epistasis to trait variation, an important phenomenon (Ehrenreich 2017) that may nonetheless explain a minority of complex trait variation in general (Bloom et al. 2015; Albert et al. 2018; Hivert et al. 2021).

### X-QTL dissection of insecticide resistance traits may benefit from the development of novel mapping populations

The DSPR is created from a set of strains derived from wild-caught individuals captured prior to widespread deployment of organophosphate insecticides (e.g., malathion), and well before routine use of many now commonly used insecticide classes (e.g., pyrethroids, neonicotinoids). And it appears the DSPR does not segregate for variation in at least one key insecticide target gene (i.e., *Ace*), variation that is known to exist in current natural populations. This suggests that a DSPR-derived base population may not be optimal for genetic dissection of all insecticide response traits via X-QTL mapping. Instead, the powerful X-QTL approach might be more profitably employed using novel, outbred, highly-recombinant multiparental populations, subject to the constraints of starting from sequenced founders to enable accurate haplotype inference from pooled sequencing with modest coverage, and allowing the founders to intercross for many generations to yield fine-scale QTL mapping. Such populations would have similar properties to a DSPR-based X-QTL design (see Macdonald et al. 2022), but enable deeper exploration of insecticide resistance in current populations. For instance, since not all the genetic variation contributing to insecticide resistance appears to takes the form of intermediate-frequency polymorphisms of large-effect (e.g., Schmidt et al. 2017; Denecke et al. 2017), and given the modest power for low-frequency and/or small-effect variants in the DGRP (Mackay and Huang 2018), an X-QTL design could be exploited to discover genes that segregate for rare or small-effect insecticide-relevant genetic variation in contemporary populations of *D. melanogaster*.

## ACKNOWLEDGEMENTS

Data collection was supported by NIH R01 OD010974 (to SJM and ADL) and NIH R01 ES029922 (to SJM), and analytical work was supported by NIH P20 GM103418. We also thank the KU Genome Sequencing Core (supported by NIH P20 GM103638).

## SUPPLEMENTARY TABLE LEGEND

**Supplementary Table 1 – Full details of all genes within mapped QTL intervals.** Presents 407 genes (344 protein-coding) and 178 genes (145 protein-coding) within the Chr2R and Chr3L mapped malathion QTL, respectively. Columns are as follows: Col 1) row number. Col 2) fbgn – FlyBase ID. Col 3) annotation_symbol. Col 4) chr – chromosome arm; since there are only 2 QTL on separate arms, this column distinguishes the QTL intervals. Col 5) gene_loc_max – the maximum (right-most) genomic position of the gene. Col 6) gene_loc_min – the minimum (left-most) genomic position of the gene. Col 7) strand – which strand (+/-) encodes the gene. Col 8) gene_name. Col 9) gene_symbol. Col 10) vocab_hit. Any gene that is a member of one of five specific FlyBase gene groups (“FBgg”) or one of two specific gene ontology (“GO”) groups is tagged with a number (1-7) indicating this. Numeric codes are: (1) FBgg0000547, ATP-BINDING CASSETTE TRANSPORTERS, (2) FBgg0001222, CYTOCHROME P450, (3) GO:0098754, detoxification, (4) FBgg0000077, GLUTATHIONE-S-TRANSFERASES, (5) FBgg0000797, GT1 FAMILY OF UDP-GLYCOSYLTRANSFERASES, (6) FBgg0001375, OTHER CARBOXYLESTERASES, (7) GO:0017085, response to insecticide. Genes not part of any of these groups are marked with a 0. Col 11) de_hit. Any gene shown to be significantly differentially-expressed in adult female gut tissue in response to malathion treatment by Salces-Ortiz et al. (2020; PMID: 32075557) is marked as such. This group independently tested 4 strains: (A) SE-Sto, (B) RAL-375, (C) RAL-377, (D) iso-1. For each strain, expression of the gene can be upregulated with treatment (“u”) or downregulated (“d”), so genes are marked as “Au”, “Ad”, “Bu”, and so on in the table. Genes not differentially-expressed in any strain are marked with 0. Note that genes determined to be “significant” in the original expression study are those that survive a strain-specific, Benjamini-Hochberg adjusted *p*-value threshold of 0.05, and show a fold-change of at least 1.5 between control and malathion treatment.

## REFERENCES

Al-Anzi, B., E. Armand, P. Nagamei, M. Olszewski, V. Sapin et al., 2010 The leucokinin pathway and its neurons regulate meal size in Drosophila. Curr Biol 20 (11):969–978.

Albert, F.W., J.S. Bloom, J. Siegel, L. Day, and L. Kruglyak, 2018 Genetics of trans-regulatory variation in gene expression. Elife 7.

Battlay, P., P.B. Leblanc, L. Green, N.R. Garud, J.M. Schmidt et al., 2018 Structural Variants and Selective Sweep Foci Contribute to Insecticide Resistance in the Drosophila Genetic Reference Panel. G3 (Bethesda) 8 (11):3489–3497.

Battlay, P., J.M. Schmidt, A. Fournier-Level, and C. Robin, 2016 Genomic and Transcriptomic Associations Identify a New Insecticide Resistance Phenotype for the Selective Sweep at the Cyp6g1 Locus of Drosophila melanogaster. G3 (Bethesda) 6 (8):2573–2581.

Bloom, J.S., I. Kotenko, M.J. Sadhu, S. Treusch, F.W. Albert et al., 2015 Genetic interactions contribute less than additive effects to quantitative trait variation in yeast. Nat Commun 6:8712.

Casida, J.E., and G.B. Quistad, 1998 Golden age of insecticide research: past, present, or future? Annu Rev Entomol 43:1–16.

Chakraborty, M., J.J. Emerson, S.J. Macdonald, and A.D. Long, 2019 Structural variants exhibit widespread allelic heterogeneity and shape variation in complex traits. Nat Commun 10 (1):4872.

Chung, H., M.R. Bogwitz, C. McCart, A. Andrianopoulos, R.H. Ffrench-Constant et al., 2007 Cis-regulatory elements in the Accord retrotransposon result in tissue-specific expression of the Drosophila melanogaster insecticide resistance gene Cyp6g1. Genetics 175 (3):1071–1077.

Clancy, D.J., and W.J. Kennington, 2001 A simple method to achieve consistent larval density in bottle cultures. Drosoph Inf Serv 84:168–169.

Cogni, R., C. Cao, J.P. Day, C. Bridson, and F.M. Jiggins, 2016 The genetic architecture of resistance to virus infection in Drosophila. Mol Ecol 25 (20):5228–5241.

Daborn, P., S. Boundy, J. Yen, B. Pittendrigh, and R. ffrench-Constant, 2001 DDT resistance in Drosophila correlates with Cyp6g1 over-expression and confers cross-resistance to the neonicotinoid imidacloprid. Mol Genet Genomics 266 (4):556–563.

Daborn, P.J., J.L. Yen, M.R. Bogwitz, G. Le Goff, E. Feil et al., 2002 A single p450 allele associated with insecticide resistance in Drosophila. Science 297 (5590):2253–2256.

Denecke, S., R. Fusetto, and P. Batterham, 2017 Describing the role of Drosophila melanogaster ABC transporters in insecticide biology using CRISPR-Cas9 knockouts. Insect Biochem Mol Biol 91:1–9.

DSPR Variant UCSC Browser, 2019, pp. Visualization of known variation segregating among DSPR founder genomes, https://goo.gl/LLpoNH.

Duneau, D., H. Sun, J. Revah, K. San Miguel, H.D. Kunerth et al., 2018 Signatures of Insecticide Selection in the Genome of Drosophila melanogaster. G3 (Bethesda) 8 (11):3469–3480.

Ehrenreich, I.M., 2017 Epistasis: Searching for Interacting Genetic Variants Using Crosses. Genetics 206 (2):531–535.

Ehrenreich, I.M., N. Torabi, Y. Jia, J. Kent, S. Martis et al., 2010 Dissection of genetically complex traits with extremely large pools of yeast segregants. Nature 464 (7291):1039–1042.

Everman, E.R., C.L. McNeil, J.L. Hackett, C.L. Bain, and S.J. Macdonald, 2019 Dissection of Complex, Fitness-Related Traits in Multiple Drosophila Mapping Populations Offers Insight into the Genetic Control of Stress Resistance. Genetics 211 (4):1449–1467.

Ffrench-Constant, R.H., P.J. Daborn, and G. Le Goff, 2004 The genetics and genomics of insecticide resistance. Trends Genet 20 (3):163–170.

Garud, N.R., P.W. Messer, E.O. Buzbas, and D.A. Petrov, 2015 Recent selective sweeps in North American Drosophila melanogaster show signatures of soft sweeps. PLoS Genet 11 (2):e1005004.

Good, R.T., L. Gramzow, P. Battlay, T. Sztal, P. Batterham et al., 2014 The molecular evolution of cytochrome P450 genes within and between drosophila species. Genome Biol Evol 6 (5):1118–1134.

Gramates, L.S., J. Agapite, H. Attrill, B.R. Calvi, M.A. Crosby et al., 2022 FlyBase: a guided tour of highlighted features. Genetics 220 (4).

Green, L., P. Battlay, A. Fournier-Level, R.T. Good, and C. Robin, 2019 Cis- and trans-acting variants contribute to survivorship in a naive Drosophila melanogaster population exposed to ryanoid insecticides. Proc Natl Acad Sci U S A 116 (21):10424–10429.

Hall, M.E., G.M. Loeb, L. Cadle-Davidson, K.J. Evans, and W.F. Wilcox, 2018 Grape Sour Rot: A Four-Way Interaction Involving the Host, Yeast, Acetic Acid Bacteria, and Insects. Phytopathology 108 (12):1429–1442.

Harrop, T.W., T. Sztal, C. Lumb, R.T. Good, P.J. Daborn et al., 2014 Evolutionary changes in gene expression, coding sequence and copy-number at the Cyp6g1 locus contribute to resistance to multiple insecticides in Drosophila. PLoS One 9 (1):e84879.

Hartley, C.J., R.D. Newcomb, R.J. Russell, C.G. Yong, J.R. Stevens et al., 2006 Amplification of DNA from preserved specimens shows blowflies were preadapted for the rapid evolution of insecticide resistance. Proc Natl Acad Sci U S A 103 (23):8757–8762.

Hawkins, N.J., C. Bass, A. Dixon, and P. Neve, 2018 The evolutionary origins of pesticide resistance. Biol Rev Camb Philos Soc.

Hemingway, J., N.J. Hawkes, L. McCarroll, and H. Ranson, 2004 The molecular basis of insecticide resistance in mosquitoes. Insect Biochem Mol Biol 34 (7):653–665.

Hemingway, J., and H. Ranson, 2000 Insecticide resistance in insect vectors of human disease. Annu Rev Entomol 45:371–391.

Hivert, V., J. Sidorenko, F. Rohart, M.E. Goddard, J. Yang et al., 2021 Estimation of non-additive genetic variance in human complex traits from a large sample of unrelated individuals. Am J Hum Genet 108 (5):786–798.

Huang, W., A. Massouras, Y. Inoue, J. Peiffer, M. Ramia et al., 2014 Natural variation in genome architecture among 205 Drosophila melanogaster Genetic Reference Panel lines. Genome Res 24 (7):1193–1208.

Karasov, T., P.W. Messer, and D.A. Petrov, 2010 Evidence that adaptation in Drosophila is not limited by mutation at single sites. PLoS Genet 6 (6):e1000924.

King, E.G., and A.D. Long, 2017 The Beavis Effect in Next-Generation Mapping Panels in Drosophila melanogaster. G3 (Bethesda) 7 (6):1643–1652.

King, E.G., S.J. Macdonald, and A.D. Long, 2012a Properties and power of the Drosophila Synthetic Population Resource for the routine dissection of complex traits. Genetics 191 (3):935–949.

King, E.G., C.M. Merkes, C.L. McNeil, S.R. Hoofer, S. Sen et al., 2012b Genetic dissection of a model complex trait using the Drosophila Synthetic Population Resource. Genome Res 22 (8):1558–1566.

King, E.G., B.J. Sanderson, C.L. McNeil, A.D. Long, and S.J. Macdonald, 2014 Genetic dissection of the Drosophila melanogaster female head transcriptome reveals widespread allelic heterogeneity. PLoS Genet 10 (5):e1004322.

Lamb, A.M., E.A. Walker, and P.J. Wittkopp, 2017 Tools and strategies for scarless allele replacement in Drosophila using CRISPR/Cas9. Fly (Austin) 11 (1):53–64.

Li, H., 2011 A statistical framework for SNP calling, mutation discovery, association mapping and population genetical parameter estimation from sequencing data. Bioinformatics 27 (21):2987–2993.

Li, H., 2013 Aligning sequence reads, clone sequences and assembly contigs with BWA-MEM. arXiv q-bio.GN.

Linder, R.A., A. Majumder, M. Chakraborty, and A. Long, 2020 Two Synthetic 18-Way Outcrossed Populations of Diploid Budding Yeast with Utility for Complex Trait Dissection. Genetics 215 (2):323–342.

Long, A.D., S.J. Macdonald, and E.G. King, 2014 Dissecting complex traits using the Drosophila Synthetic Population Resource. Trends Genet 30 (11):488–495.

Macdonald, S.J., K.M. Cloud-Richardson, D.J. Sims-West, and A.D. Long, 2022 Powerful, efficient QTL mapping in Drosophila melanogaster using bulked phenotyping and pooled sequencing. Genetics 220 (3).

Mackay, T.F., S. Richards, E.A. Stone, A. Barbadilla, J.F. Ayroles et al., 2012 The Drosophila melanogaster Genetic Reference Panel. Nature 482 (7384):173–178.

Mackay, T.F.C., and W. Huang, 2018 Charting the genotype-phenotype map: lessons from the Drosophila melanogaster Genetic Reference Panel. Wiley Interdiscip Rev Dev Biol 7 (1).

Magwire, M.M., D.K. Fabian, H. Schweyen, C. Cao, B. Longdon et al., 2012 Genome-wide association studies reveal a simple genetic basis of resistance to naturally coevolving viruses in Drosophila melanogaster. PLoS Genet 8 (11):e1003057.

McDonnell, C.M., D. King, J.M. Comeron, H. Li, W. Sun et al., 2012 Evolutionary toxicogenomics: diversification of the Cyp12d1 and Cyp12d3 genes in Drosophila species. J Mol Evol 74 (5-6):281–296.

Menozzi, P., M.A. Shi, A. Lougarre, Z.H. Tang, and D. Fournier, 2004 Mutations of acetylcholinesterase which confer insecticide resistance in Drosophila melanogaster populations. BMC Evol Biol 4:4.

Moyes, C.L., J. Vontas, A.J. Martins, L.C. Ng, S.Y. Koou et al., 2017 Contemporary status of insecticide resistance in the major Aedes vectors of arboviruses infecting humans. PLoS Negl Trop Dis 11 (7):e0005625.

Mutero, A., M. Pralavorio, J.M. Bride, and D. Fournier, 1994 Resistance-associated point mutations in insecticide-insensitive acetylcholinesterase. Proc Natl Acad Sci U S A 91 (13):5922–5926.

Najarro, M.A., J.L. Hackett, and S.J. Macdonald, 2017 Loci Contributing to Boric Acid Toxicity in Two Reference Populations of Drosophila melanogaster. G3 (Bethesda) 7 (6):1631–1641.

Najarro, M.A., J.L. Hackett, B.R. Smith, C.A. Highfill, E.G. King et al., 2015 Identifying Loci Contributing to Natural Variation in Xenobiotic Resistance in Drosophila. PLoS Genet 11 (11):e1005663.

Rivero, A., J. Vezilier, M. Weill, A.F. Read, and S. Gandon, 2010 Insecticide control of vector-borne diseases: when is insecticide resistance a problem? PLoS Pathog 6 (8):e1001000.

Salces-Ortiz, J., C. Vargas-Chavez, L. Guio, G.E. Rech, and J. Gonzalez, 2020 Transposable elements contribute to the genomic response to insecticides in Drosophila melanogaster. Philos Trans R Soc Lond B Biol Sci 375 (1795):20190341.

Schmidt, J.M., P. Battlay, R.S. Gledhill-Smith, R.T. Good, C. Lumb et al., 2017 Insights into DDT Resistance from the Drosophila melanogaster Genetic Reference Panel. Genetics 207 (3):1181–1193.

Schmidt, J.M., R.T. Good, B. Appleton, J. Sherrard, G.C. Raymant et al., 2010 Copy number variation and transposable elements feature in recent, ongoing adaptation at the Cyp6g1 locus. PLoS Genet 6 (6):e1000998.

Schrider, D.R., D.J. Begun, and M.W. Hahn, 2013 Detecting highly differentiated copy-number variants from pooled population sequencing. Pac Symp Biocomput:344–355.

Scott, J.G., and N. Buchon, 2019 Drosophila melanogaster as a powerful tool for studying insect toxicology. Pestic Biochem Physiol 161:95–103.

Shi, M.A., A. Lougarre, C. Alies, I. Fremaux, Z.H. Tang et al., 2004 Acetylcholinesterase alterations reveal the fitness cost of mutations conferring insecticide resistance. BMC Evol Biol 4:5.

Soetaert, K., K. Van den Meersche, and D. van Oevelen, 2009 Package limSolve, Solving Linear Inverse Models in R.

Spencer, C.C., Z. Su, P. Donnelly, and J. Marchini, 2009 Designing genome-wide association studies: sample size, power, imputation, and the choice of genotyping chip. PLoS Genet 5 (5):e1000477.

Steinmetz, L.M., H. Sinha, D.R. Richards, J.I. Spiegelman, P.J. Oefner et al., 2002 Dissecting the architecture of a quantitative trait locus in yeast. Nature 416 (6878):326–330.

Stern, D.L., 2014 Identification of loci that cause phenotypic variation in diverse species with the reciprocal hemizygosity test. Trends Genet 30 (12):547–554.

Sun, H., N. Buchon, and J.G. Scott, 2017 Mdr65 decreases toxicity of multiple insecticides in Drosophila melanogaster. Insect Biochem Mol Biol 89:11–16.

Tudi, M., H. Daniel Ruan, L. Wang, J. Lyu, R. Sadler et al., 2021 Agriculture Development, Pesticide Application and Its Impact on the Environment. Int J Environ Res Public Health 18 (3).

World Health Organization, 2006 Pesticides and their application, for the control of vectors and pests of public health importance (WHO/CDS/NTD/WHOPES/GCDPP/2006.1).

Xu, C., C.Y. Li, and A.N. Kong, 2005 Induction of phase I, II and III drug metabolism/transport by xenobiotics. Arch Pharm Res 28 (3):249–268.

